# LAMBDR: Long-range amplification and Nanopore sequencing of the *Mycobacterium bovis direct-repeat region*. A novel method for in-silico spoligotyping of *M. bovis* directly from badger faeces

**DOI:** 10.1101/791129

**Authors:** R.S. James, E.R. Travis, A. D. Millard, P.C. Hewlett, L. Kravar-Garde, E.M. Wellington

## Abstract

The environment is an overlooked source of *Mycobacterium bovis*, the causative agent of bovine TB. Long read, end to end sequencing of variable repeat regions across the *M. bovis* genome was evaluated as a method of acquiring rapid strain level resolution directly from environmental samples. Eight samples of *M. bovis*, two BCG strains (Danish and Pasteur), and a single *M. tuberculosis* type culture (NCTC 13144) were used to generate data for this method. Long range PCR amplification of the direct repeat region was used to synthesize ∼5kb template DNA for onward sequence analysis. This has permitted culture independent identification of *M. bovis* spoligotypes present in the environment. Sequence level analysis of the direct repeat region showed that spoligotyping may underestimate strain diversity due to the inability to identify both SNPs and primer binding mutations using a biotinylated hybridisation approach.

## Introduction

Sequencing pathogens directly from the environment allows rapid epidemiological studies to be undertaken in response to outbreaks of disease. Bovine tuberculosis (bTB) is a progressive pulmonary disease of the bovidae that remains endemic to many cattle populations around the world. *Mycobacterium bovis*, the causal agent of bTB, can persist in the environment and this may contribute to further infection in domesticated livestock and wildlife (Courtenay et al., 2006, King et al., 2015). Infected animals shed to the environment via aerosol, urine and faeces, with a growing body of evidence suggesting that environmental *M. bovis* could play an important role in the persistence of this disease (Wellington and Courtenay, 2014, Duffield and Young, 1985, King et al., 2015, Barbier et al., 2017). Previous studies report the molecular detection of *M. bovis* in environmental faecal samples from the European badger (*Meles meles*) upwards of 15 months after excretion (Young et al., 2005). However, the isolation and culture of *M. bovis* is rarely successful from environmental samples due to the difficulty in selectively isolating *M. bovis* from diverse and competitive microbial communities. While the molecular detection of *M. bovis* in environmental samples has provided a useful marker for tracking diseased populations (King et al., 2015), information relating to strain type diversity could only previously be obtained by culturing isolates from infected individuals. Here we present a new method to strain type *M. bovis* directly from environmental samples, such as badger faeces, with the aim to better understand the role of the environment in the epidemiology of bovine tuberculosis.

The roles of wildlife reservoirs have become synonymous with the spread of bTB throughout the world. Animals such as the white-tailed deer (*Odocoileus virginianus*), common brush tail possum (*Trichosurus Vulpecula)*, wild boar (*Sus scrofa*) and European badger (*Meles meles*) have all been shown to be susceptible to this disease and maintain an infective load at the population level (O’Brien et al., 2002, Nugent, 2011, Martin-Hernando et al., 2007, Fitzgerald and Kaneene, 2012). Badgers are an important wildlife reservoir of *M. bovis* in the United Kingdom (Donnelly et al., 2003) and infected badgers have been shown to shed *M. bovis* in their faeces (King et al., 2015). Social groups of badgers dig underground tunnel systems known as setts and defecate into communal “latrines” which are often located on cattle pasture.

*M. bovis* is a highly genetically homogeneous species. For example, 98.9% of the *M. bovis* genome is conserved between strains with the 16S rRNA gene fully conserved within the *Mycobacterium tuberculosis* complex (MTC). However, whole genome sequencing of *M. bovis* directly from environmental samples is technically challenging and often limited by the low abundance of the organism in the sample type. The use of the direct repeat region and variable nucleotide tandem repeats (VNTRs) have been reported to act as robust proxies for strain type diversity (Brudey et al., 2006, Roring et al., 2002, Zeng et al., 2016, Barbier et al., 2016). The direct repeat region of the MTC is a ∼5 kb region of the genome belonging to the CRISPA sequence family. This region is comprised of repeated subunits interspersed with spacer DNA. The presence or absence of the 43 different spacer DNA subunits can be used to fingerprint and identify divergent lineages of *M. bovis* and *M. tuberculosis* using a biotinylated amplification and hybridization approach. A limitation to this method is that it relies on culture from an infected individual and requires multiple PCR reactions per sample that are likely to be subject to inhibition when directed at environmental samples such as soil and faeces (Pontiroli et al., 2011).

The ability to detect and type members of the mycobacterium complex, such as *M. bovis* and *M. tuberculosis*, without the requirement of culture provides a rapid and low cost epidemiological tool that can be implemented in countries where *M. bovis* and *M. tuberculosis* are endemic and zoonotic pathogens to both humans and livestock. This is of particular relevance to LMIC countries where diagnostic costs are high and co-infections are present. In this study we developed long range PCR primers to amplify the direct repeat region of the *M. bovis* genome and then used the portable Oxford nanopore MinION to undertake long-read, end to end amplicon sequencing. Data is presented on the performance of this assay on pure culture isolates of BCG, wild type *M. bovis isolates, M. tuberculosis* type culture, inoculated environmental samples and naturally infected badger faeces. This has allowed us to differentiate known strain types from both culture and the environment.

## Methods

### Sample collection

*Mycobacterium bovis* variant BCG NCTC 5692 and NCTC 14044, *M. bovis* NCTC 10772 and *M. tuberculosis* type strain NCTC 13144 were sourced from the Public Health England culture collection. Six wildtype *M. bovis* heat treated lysates were also supplied by Public Health England, originating from infected livestock from the south west of England. Badger faecal samples were collected from Woodchester Park, Gloucestershire, UK. Negative badger faeces used for spiking and negative controls were sourced from APHA captive animals. Samples were stored at −20 ° C prior to DNA extraction. A water sample from a farmland drinking trough contaminated with low levels of *M. bovis* was also included in the development of this method.

Six samples of known negative badger faeces inoculated with tenfold serial dilutions of BCG. These dilutions ranged from 1 × 10^8^ genomic equivalents g^-1^ of faeces to 1 × 10^3^ genomic g^-1^. Samples were homogenised via stirring with a sterile pipette tip for one min and frozen at – 20 ° C prior to DNA extraction.

### DNA extraction

All DNA was extracted using the MolBio DNA fast DNA extraction kit for soil (cat no: 6560-200) as per manufacturers instruction with a modifications (Sweeney et al). Silica based binding matrix beads were suspended and washed twice, once with the supplied wash buffer and once with 80 % EtOH. DNA was eluted in 50 μl of warmed DES elution buffer at 56 ° C and quantified using a Qbit high sensitivity assay.

### PCR amplification of the direct repeat region

Three PCR primer sets for the amplification of the 5 kb direct repeat region were computed using PRIMER 3 (Table 1). Estimated annealing temperatures both within and between primers were maintained within +-0.5 ° C. Efforts were made to limit cross species homology outside of the MTC using the NCBI database. Maximum permissible homopolymer sites were reduced to 3 and self-compatibility was limited to less than 2 at the 3’ end. Primer tertiary structures and primer dimer formation were analysed using Oligoanalyzer. No conformational tertiary structures or primer dimers were permitted with a delta G greater than −6. Primer sequences are shown in Table 1.

**Table 1.**
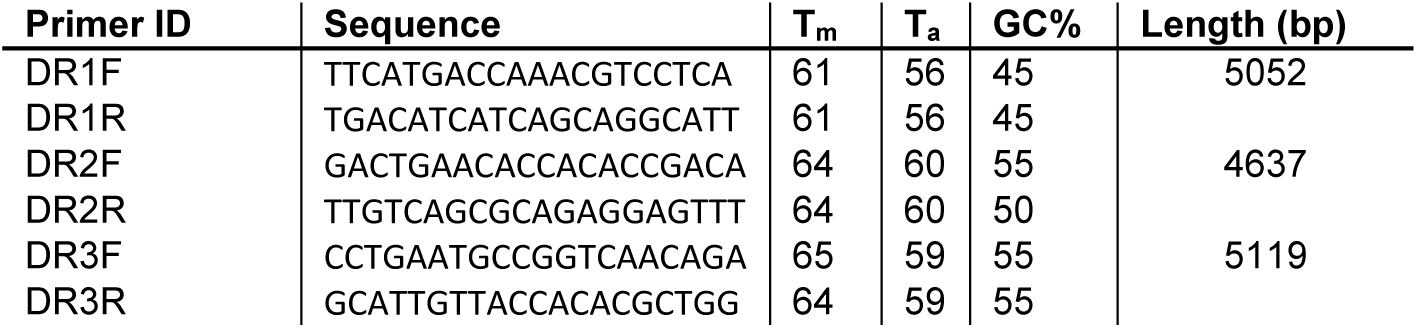
Primer sequence, annealing temperature and amplicon size tested in this study.

One 25 μl PCR reaction consisted of 1 μl of NEB hotstart long amp Taq (Cat no:M0534L), 5 μl of NEB Long amp Taq buffer, 300μM dNTP mix, 10 μg/ul BSA, 200 ng NEB EtSSB, 0.4 μM forward and reverse primers, and 5 μl of DNA template. PCR was undertaken using an Eppendorf Master cycler under the conditions 94 ° C for 2 min then 35 cycles of 30 sec at 94 ° C, 45 sec at 58 ° C and 5 min at 65 ° C. A final extension phase of 10 min at 65 ° C was used. PCR products were then exposed to 50 ng of Proteinase K and digested at 56 ° C for five min. Products then underwent a 0.4 x SPRI clean up using 80% EtOH as a wash buffer. Samples were eluted into 50 μl of molecular grade H_2_O at 37 ° C for 30 min. DNA was standardised to 1.5 μg in 50 μl of H_2_O and then stored at −20 ° C prior to onward analysis.

### PCR amplification of the direct repeat region

Amplification of the direct repeat region from *M. bovis BCG, M. bovis* and *M. tuberculosis* was undertaken using the three potential primer sets under gradient PCR conditions of T_a_ of 54.5 ° C : 65 ° C using an Eppendorph Mastercycler. Amplification of the direct repeat region from the six wildtype isolates, spiked faecal samples and a naturally infected badger faecal sample was undertaken using primer set one using end-point PCR. Off-target PCR amplification was assessed using pure culture DNA from *M. avium, M. intracellulare, M. abscessus, and M. fortuitum*.

### Quantification of genomic copy number

The number of genomic copies of *M. bovis* and *M. bovis* BCG present in both spiked and naturally infected samples were quantified using Taq-man qPCR assay as described in King et al. (2015).

### Library preparation

Library preparation was undertaken using the Oxford Nanopore SQK-LSK 109 ligation sequencing kit and native barcode kit NB003. End repair, dA tailing and FFPE repair was undertaken in parallel using 48 μl of PCR product standardised to 1.5 ug. DNA was eluted into 12 μl of H_2_O. Sequencing was undertaken on a MinION using R 9.4.1 flow cells (FLO-MIN 106) and MinKNOW version 2.0.1. Two multiplexed libraries were prepared, one consisting of amplicons generated from *M. bovis* BCG pasture, BCG Danish, *M. bovis* and *M. tuberculosis* cultured isolates and one consisting of *M. bovis* BCG Danish, six wildtype isolates, one artificially spiked badger faeces with BCG Danish at 1 × 10^6^ copies g^-1^ one naturally infected badger faeces at 1 × 10^8^ copies g^-1^. Libraries were run independently on two different flowcells (R 9.4.1). Once sequencing was completed, fast5 reads were basecalled using Guppy 2.0. using the high accuracy configuration. Reads with a Q score of < 8 were discarded.

### Sequence assembly

Fastq reads were demultiplexed by barcode identity using qCAT. Reads with middle adaptors detected were discarded from this analysis. Reads were then mapped to the direct repeat region of reference genomes NC_000962.3, AM408590.1 and GCF_000195835.2 using minimap2 and extracted using Samtools. Reads were randomly down sampled to 1000 reads using fastqSample with reads < 3.5 kb discarded. Contigs were assembled using the program Canu V 1.8. Fastq reads were then mapped back to the consensus sequences using minimap2 and polished using Nanopolish v 0.11. Spoligotype analysis was then undertaken using Spo-Typing v2.0 (Xia et al., 2016). Strain fingerprints were then compared with the national *M. bovis* spoligotype database, worldwide TB strain database and NCBI reference genomes. Sequence level homology to known BCG type strains were assessed using MUMmer v 3.2.3 and Nucmer (Kurtz et al., 2004). Sequence level homology between isolates was assessed using a MUSCLE alignment with reduced gap open and gap extension penalties. Sequence alignments were analysed in a Maximum likelihood tree using MEGA X.

## Results

### PCR amplification of the direct repeat region from pure culture

All three primer sets for amplification of the 5kb direct repeat region showed strong amplification when characterised using pure culture DNA from *M. bovis* BCG (Danish) M. bovis BCG (Pasteur), *M. bovis* and *M. tuberculosis*. Primer set one was chosen for further analysis due to the strong amplification and homology of primer annealing temperatures.

### Amplification of the direct repeat region in environmental samples

Six wild type isolates showed positive amplification when using primer set one. Spiked badger DNA templates showed amplification at 1 × 10^4^ copies g^-1^. Amplification of the direct repeat region was also achieved in a naturally infected badger faeces quantified using qPCR at 1 × 10 ^8^ copies g^-1^. Low level amplification was observed in contaminated drinking trough water with some off target amplification. No off-target amplification was seen for *M. abscesses, M. avium*, or *M. fortuitum*. Low level amplification of a 2kb fragment was seen in response to *M. intracellulare*.

### Sequencing and assembly

The two sequencing reactions were run for two hours generating approximately 1.54 gb and 1.23 gb of data in 542041 and 461010 reads respectively. An N50 value for each run was calculated at 4216 bp and 3892 bp respectively prior to demultiplexing and quality control. Assembled contig length, coverage, N50 and reference sequence homology for each sample are shown in table 2. Only partial sequencing of the direct repeat region was achieved from contaminated drinking trough water.

**Table 2.**
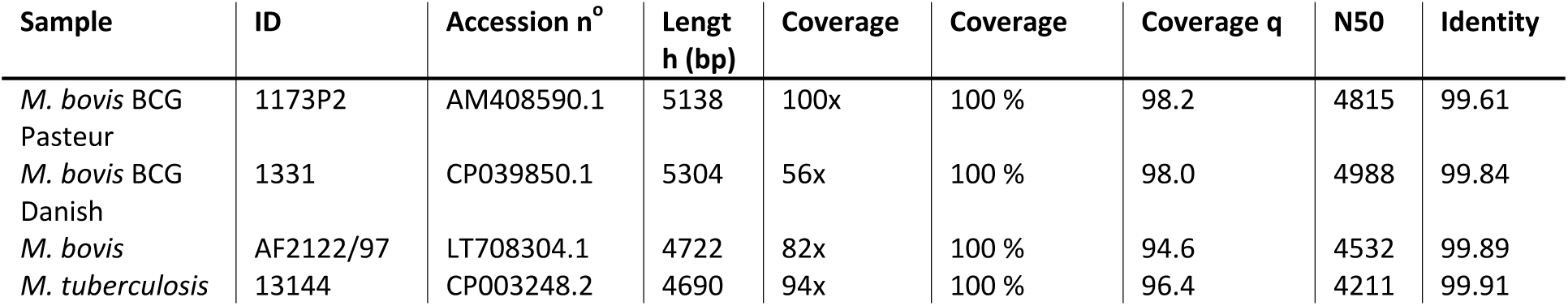
Assembled and polished contig length, coverage, N50 and reference sequence homology for each type sample sequenced in this study. Reference sequence identity calculated using nucmer.

### In silico spoligotypes analysis

Successful spoligotypes were generated for all samples using Spo-Type V 2.0 (Table 3). Spoligotypes for all non BCG strains successfully matched to existing databases (Table 3). Multiple sequence alignments clustered *M. bovis* BCG strains with known reference genomes and clustered separately from Wild Type strains of *M. bovis* (Figure 1).

**Table 3.**
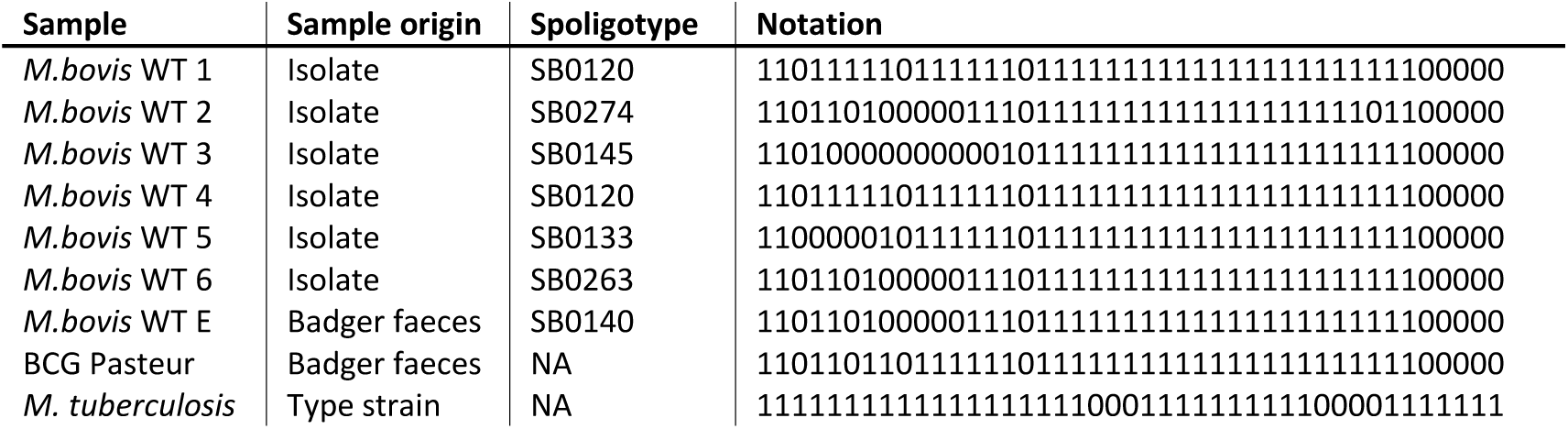
In-silico spoligotypes, generated for each wild type sample tested in this study. Spoligotypes were generated automatically using Spoligotyper-v2.0.

**Figure 1.**
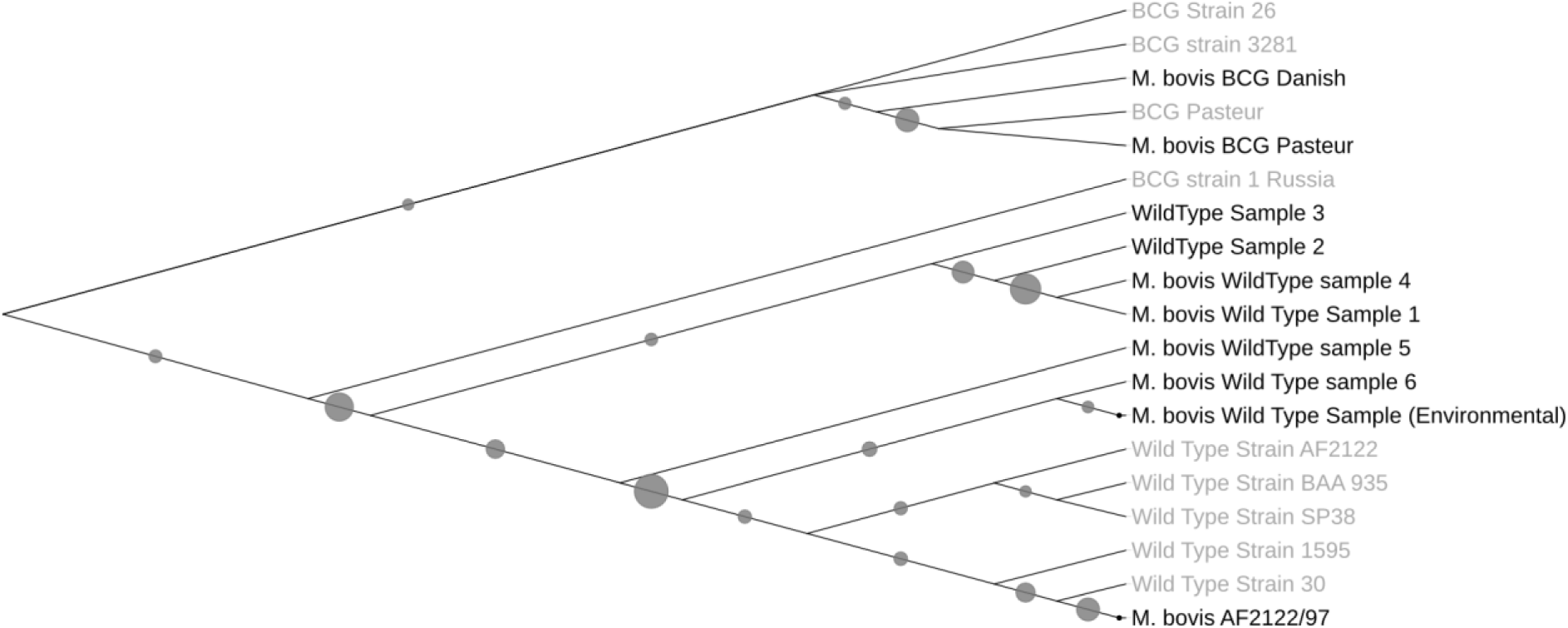
Clustering of *M. bovis* strains using spoligotyping amplicon sequence level diversity. A maximum likelihood tree (bootstrap n = 500) based on sequence similarity of the five kb direct repeat region. Clustering is relative to Genbank reference sequences shown in grey, bootstrap values are denoted by circles.

## Discussion

*In silico* spoligotyping was successfully achieved for both *M. bovis* and *M. tuberculosis* using pure cultures. Furthermore, in silico spoligotyping was also achieved with DNA extracted from *M. bovis* isolates, spiked faecal samples and naturally infected badger faeces. Partial sequencing was achieved for the direct repeat region from contaminated drinking trough water. While previous studies have shown it possible to undertake whole genome sequencing of pure isolates as well as partial sequencing and assembly of *M. tuberculosis* from low complexity human sputum samples (George et al., 2018), this is the first study to present a feasible method for resolving a degree of strain level differentiation directly from environmental reservoirs of infection. It is likely that the incomplete spoligotyping of contaminated drinking tough water is due to the low level of M. bovis within the sample which indicates that this strain typing tool is less sensitive than the qPCR method to identify and quantify *M. bovis* in environmental samples (King et al., 2015, Courtenay et al., 2006). However, this method is suitable for rapid, high throughput, low-cost in-house spoligotyping of both *M. bovis* at 1 × 10^4^ copies g^-1^ in badger faeces and *M. tuberculosis* from culture. The ability to differentiate *M. bovis* from *M. tuberculosis* based on the presence of the final five unique spacer DNA sequences in the direct repeat region (Niemann et al., 2000) also highlights the application of this method in regions where *M. bovis* and *M. tuberculosis* are present within both the human and wildlife population.

PCR primers developed for the amplification of variable regions of the *M. bovis* and *M. tuberculosis* genome were shown to amplify target regions of DNA from pure culture, spiked faecal samples and a naturally infected badger faecal sample. The lack of off-target and non-specific amplification in closely related species from the *M. tuberculosis* complex indicates that this method is suitable for specific amplification of target DNA in some complex environmental matrices. However, due to the partial amplification of off target DNA in samples containing low levels of M. bovis, pre-processing clean up and post processing bioinformatics is recommended to map reads to a known reference data base before assembly.

Amplification of target DNA from spiked environmental samples was achievable down to 1 × 10 ^4^ genomic equivalents g^-1^. This is likely due to the presence of PCR inhibitors present in faecal samples and the mechanical method of DNA extraction required to recover DNA from resilient bacteria such as *M. bovis*. While total fragment length of the DNA extract was approximated at between 5 - 10 kb using gel electrophoresis, the effects of DNA fragmentation in low copy number infected samples has not been quantified in this study. This may explain, in part, the reduction in sensitivity of this assay when used in low abundance samples as recovery of full-length template DNA may be limited under these conditions. Despite this shortfall we have shown that amplification of the direct repeat region from positive wild badger faeces with a realistic infective load (King et al., 2015) is achievable using this extraction method.

Using the direct repeat region for in-silico spoligotyping gives a degree of resolution comparable with existing methods. The ability to inspect the sequence level resolution of these variable regions increases the potential resolution and reduces the PCR based artefacts associated with the spoligotyping technique. While it is difficult to quantify the accuracy of sequence level resolution when analysing uncharacterised samples, our comparison of known samples of BCG to reference genomes indicates a > 99.8% homology is achievable using polished contigs. Sequence level phylogenetic analysis of all the sequences in this data set also indicates that the distinct lineages of *M. bovis, M. tuberculosis* and *M. bovis BCG* may be possible with larger data sets.

The data presented provides evidence of a suitable method to spoligotypes wild type *M. bovis* present in badger faeces in the UK. Furthermore, this method is shows the potentially resolve *M. tuberculosis* spoligotypes from non-invasive sampling methods such as neonatal faeces and sputum. We feel this is a valid contribution to the global epidemiology of both *M. bovis* and *M. tuberculosis* as it offers a high through-put and low cost alternative to isolation and culture. While the amount of sequence data recovered for each isolate in this study far surpassed the requirements for accurate sequence assembly and coverage in the majority of samples, future applications using smaller capacity flow cells are likely to substantially further reduce costs of this application without sacrificing sequence level accuracy.

